# Hypothesis-Testing Improves the Predicted Reliability of Neuroscience Research

**DOI:** 10.1101/537365

**Authors:** Bradley E. Alger

## Abstract

Critics often cite statistical problems as prime contributors to the “reproducibility crisis” of science, expressing great concern about research that bases major conclusions on single p-valued statistical tests. The critics also argue that the predicted reliability of neuroscience research in particular is low because much of the work depends heavily on small experimental sample sizes and, hence, its statistical tests lack adequate “power.”

It isn’t known how common the practice of basing major conclusions on single tests is in neuroscience or how the statistical criticisms affect the validity of conclusions drawn by laboratory research that evaluates hypotheses via multiple tests. I review a sample of neuroscience publications to estimate the prevalence and extensiveness of hypothesis-testing research. I then apply R.A. Fisher’s method for combining test results to show that the practice of testing multiple predictions of hypotheses increases the predicted reliability of neuroscience research.

## Introduction

In recent years the reliability of science has been called into question and concerns raised that science is suffering from a “reproducibility crisis.” Indeed, the statistician, John Ioannidis, has argued that “most published research findings are false,” especially in biomedical science (1). However, biomedical science encompasses many experimental approaches and not all may be equally susceptible to reproducibility concerns tied to statistical practices. For instance, Ioannidis suggests

“… that the high rate of nonreplication (lack of confirmation) of research discoveries is a consequence of the convenient, yet ill-founded strategy of *claiming conclusive research findings solely on the basis of a single study assessed by formal statistical significance, typically for a p-value less than 0.05***.”** [Italics added.] He continues, **“**Research is not most appropriately represented and summarized by *p*-values, but, unfortunately, there is a widespread notion that medical research articles should be interpreted based only on *p*-values.”

His comments leave open the possibility that science in which single, p-valued significance tests are not accepted decisive evidence would be more reliable than science in which they are. On the other hand, statisticians also argue that much experimental science, such as neuroscience, is fatally flawed because its claims are based on statistical tests that are “underpowered,” largely because of small experimental group sizes (2). The statistical power of a test is usually designated “1-β,” where β is the probability of failing to reject the null hypothesis when it should be rejected (i.e., the chance of missing a true difference between groups). Statistical power varies from 0 – 1 and values of ≥ 0.8 are considered “good”. Button et al. (2), calculate that the typical power of a neuroscience study is 0.2, i.e., quite low.

The likelihood of reproducing a scientific result is frequently estimated by a parameter called Positive Predictive Value (PPV), defined as “the post-study probability that [the experimental result] is true” (1). In addition to the “pre-study odds” of a result’s being correct, the PPV (formula given in equation 2 below) is heavily dependent on the p-value of the result and the statistical power of the test. The PPVs estimated for neuroscience research are low (2).

In summary, the statisticians’ criticisms of neuroscience research rest on the unproven assumptions that: 1) neuroscience findings are determined by single experimental tests and 2) the reliability of neuroscience conclusions is necessarily undermined by the use of small experimental group sizes. To evaluate the first assumption, I analyzed papers published in a leading journal of neuroscience. To evaluate the second assumption, I review a method devised by R.A. Fisher (the Fisher Method for Combining Results) for putting together the p-values of several tests of a single hypothesis and calculating the probability of obtaining the aggregate result of all the tests (3,4). Finally, I show that the probability estimate derived from the Fisher Method leads to estimates of PPV that are much higher than previously derived.

## Methods

To gauge the applicability of the statistical criticisms to typical neuroscience research, I examined all of the Research Articles (i.e., no Brief Communications or other forms of publication) that appeared in the first three issues of *The Journal of Neuroscience* in 2018. The goals were first, to find out how many papers based major conclusions on the outcome of a single p-valued test and, second, to find out what kinds of approaches were represented. It is universally accpeted that there are different “modes” of doing science; e.g., “hypothesis-testing,” “questioning,” “Discovery,” etc., that have different standards for acceptable evidence, for decision-making, and other aspects. Experience had suggested that hypothesis-testing would be a major mode and, therefore, I began with a pdf search of each article for “hypoth” (to catch all forms referring to scientific hypothesis and avoid “statistical hypothesis” as well as instances where “hypothesis” was improperly used as a synonym for “prediction”), “predict” and “model” (counted when used synonymously with “hypothesis;” I omitted “animal models,” “model systems,” etc.) and checked every occurrence of these key words in the text to see how it was actually used. This search proved to be only marginally useful, however, as I’ll explain. After the search, I read the Abstract and Significance Statement of each paper and as much of the Introduction, Results, and Discussion as necessary to understand its objective as well as the experiments done to reach and justify its conclusion(s). I classified each paper as representing “hypothesis-based,” “Discovery science,” (identifying and characterizing the elements of a field), “Open-ended questioning,” or “computational-modeling.” There were no examples of Big Data analytics, another potentidal mode.

In identifying papers as hypothesis-based or non-hypothesis-based, I looked not only at what the authors said about their investigation – i.e. whether they stated explicitly that they were testing a hypothesis or not – but what they actually did. The papers were remarkably inconsistent in their use of the term “hypothesis,” whether it appeared at all, and what was meant by it (which is why the pdf search wasn’t very valuable). Perhaps as a result of the inconsistency, the word was often omitted entirely, even when the investigators were obviously testing hypotheses. Hence, I distinguished between “explicit” hypothesis testing, where the authors stated fairly clearly their main hypothesis, and “implicit” hypothesis testing where the authors did not use the word, “hypothesis” despite the fact that they were obviously testing one. For example, when the authors implied that some phenomenon had a potential *explanation*, then conducted a series of experimental tests of *predictions* of that explanation, and drew a final *conclusion* that reflected back on the likely validity of the original explanation, I counted it as an “implicit” hypothesis-based paper, even if the words “hypothesis,” “prediction,” etc., never appeared. For all hypothesis-testing papers, I counted the number of experimental manipulations that tested the main hypothesis. If, on the other hand, a paper pursued a series of questions or issues, but did not actually test predictions of a potential explanation – meaning that the experiments could not falsify a hypothesis no matter what their outcomes – then I categorized it as “question-based” or “Discovery” science.

While my classifications were unavoidably subjective, I think that the majority would be uncontroversial. Since I did not study the experimental findings of all papers in detail and, of course, am not an expert in all of the scientific areas covered, I undoubtedly missed nuances of the reports. Thus, for example, the counts of the numbers of independent tests of each hypothesis are probably low estimates.

## Results

I analyzed all the Research Articles in the first three issues of *The Journal of Neuroscience* in the year 2018 (n = 52). As the official journal of the Society for Neuroscience that publishes papers in all subdisciplines of neuroscience, it should be reasonably representative of the field. I classified 42 of the 52 as hypothesis-based, with 20 “explicitly” testing a hypothesis (or two, and in one case, three hypotheses) and 22 “implicitly” testing one or more hypotheses (see Supplemental Figure 1 and Methods for descriptions). Of the remaining 10 papers, 7 were “question” or “discovery” based and 3 were primarily computer-modelling studies that included a few experiments. Because the premises underlying the non-hypothesis testing kinds of studies are fundamentally distinct from hypothesis-testing studies, the same standards cannot be used to evaluate them and I did not consider papers further.

First, I noted that none of the papers, in any mode, based its conclusions on a single test. Indeed, I estimate that the summary conclusion of each hypothesis-based investigation was supported by ~7 experiments (7.5 ± 2.26, mean ± s.d., n = 42; this number does not include independent tests of alternative or secondary hypotheses). In most cases (30/42) the experimental tests were exclusively “consistent” with the overall hypothesis, but in 11/42 papers a significant portion of the reported work explicitly falsified either the main hypothesis or an expressed alternative hypothesis. In all studies, the investigators tested multiple predictions of their primary hypothesis in reaching their conclusions (Supplemental Table 1).

Theoretical statistical arguments imply that focused hypothesis-testing studies should be much more reliable than, e.g., large-open ended experimental analyses (1,2). Since most (81 %) of the neuroscience papers were based on hypothesis-testing, on theoretical grounds alone, this form of neuroscience research should be reasonably reliable. On the other hand, most of the experiments in the hypothesis-based papers were conducted on small groups of subjects, either laboratory animals or humans, and it is doubtful that they would meet rigorous criteria for adequate statistical power; a fact which should detract from their reliability. Later I will suggest how the tension between these two opposing viewpoints might be resolved.

Despite their reliance on underpowered experiments, intuitively we expect conclusions bolstered by several lines of evidence to be more secure than those resting on a single piece of evidence. Can we quantify this intuition?

Consider the analogy of tossing a fair coin 4 times; even though the chance of getting a head on any given trial is 1/2, the chance of getting a sequence of 4 heads in a row is the product of the individual probabilities, (1/2)^4^ = 1/16. In this case, each toss is a Bernoulli trial, where there are only two outcomes and each outcome is random and independent of the other trials. If we test a hypothesis with several independent tests of its predictions, we need an analogous way of combining the results and getting a value for the probability of obtaining all results. However, experimental laboratory tests are not Bernoulli trials and we cannot use the product of the p-values to evaluate the collection of test outcomes.

Fortunately, in 1939 R.A. Fisher reported the following formula for aggregating p-values for exactly this purpose:

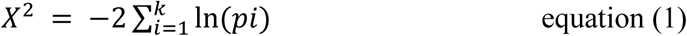

where p_i_ is the p-value of the i^th^ test and there are *k* tests in all (3,4). The sum the natural logarithms (*ln*) of the p-values multipled by −2 is a chi-squared variable with 2*k* degrees of freedom which is evaluated via a table of critical values for the chi-squared distribution (e.g., https://en.wikibooks.org/wiki/Engineering_Tables/Chi-Squared_Distibution). The table gives the probability of obtaining the cluster of all results with that chi-squared value.

Effectively, Fisher’s Method evaluates the probability of the aggregate effect by positing that there is a “global null” hypothesis that is formed by the intersection of the individual test set distributions. Rejection of the global null implies that none of the constituent null hypotheses was true; failure to reject the global null implies that the null hypothesis was true for at least one of the constituent tests.

As an example, if you have a single test result that is significant at p = 0.05, the chance of wrongly rejecting the null hypothesis and concluding that the result is significant is 1/20. But if you tested four predictions of a hypothesis at the same significance level then, according to Fisher’s Method, the probability of rejecting the null hypothesis for all four results by chance alone would be < 0.005, i.e., less than 1/200. You’d feel more confident in making decisions based on the group of results.

With Fisher’s Method you use the actual probability of each experimental outome, e.g. p = 0.041, rather than an *a priori* significance level. And, indeed, Fisher’s Method does not assume a particular significance level for each test; e.g., if the p-value of a test outcome were 0.072, then you use p = 0.072 in the calculation of the combined value. Using actual p-values like this may have advantages for scientific practice (see Discussion).

A frequent concern in the debate about reproducibility is that the predicted reliability of a scientific study, as quantified by its Positive Predictive Value (PPV) is often low. PPV is quantified as follows:

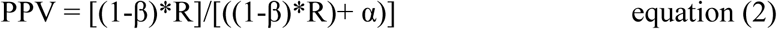

where R represents the “pre-study odds” that a hypothesis is correct, α is the p-value, and 1-β is the power of the test. R represents the anticipated number of true (T) hypotheses divided by the total number of hypotheses being evaluated, True plus False; i.e.,. R = T/(T+F). Ioannidis (1) gives an example like this: consider a “gene-screen” experiment that evaluates 1000 genes, i.e.,1000 distinct “hypotheses” where only one gene is expected to be the “correct” one. In this case, R is ~1/1000 and, with a p-value for evaluating each candidate gene of 0.05, even presuming good statistical power (≥ 0.8), PPV would be quite low, ~ 0.01. That is, the result would have roughly a one-in-a-hundred chance of being replicated, which supports the conclusion that a great deal of science is irreproducible.

However, if previous experimental work had reduced the number of alternative hypotheses to only a few, as is often the case in laboratory studies, R will be relatively high and PPV will also be higher. For example, if there were only three alternative hypotheses (it was rare to find so many in the literature that I reviewed) then R would be 1/3; i.e., ~333 times greater than the gene-screen example. Even assuming the low statistical power of ~ 0.2 that Button et al. say characterizes much neuroscience research (2) and a p-value of 0.05, PPV would be ~0.57, which implies that experimental science that evaluates a limited set of hypotheses would, in this example, be over 50 times more likely to be reliable than the open-ended gene-screen study. Yet, although this would represent a major improvement, a PPV of 0.57 would mean that the expected reproducibility of the result would be just slightly better than 50-50.

The problem is that, despite the gain in reliability expected of narrowly focused, hypothesis-based studies, the PPV calculation is still founded on the dubious assumption that experimental results are well-characterized by a single, modest p-value. Since the likely validity of an overall conclusion of an investigation is more important than that of any single result, I suggest that the calculation of PPV should reflect the aggregate probability of getting the cluster of results that all point to a given conclusion. In other words, we should use the probability obtained by the Fisher Method, p_FM_, in place of α, to calculate PPV. Making this change should have a major effect on predicted reliability, since p_FM_ will generally be lower than α and, as equation 2 shows, PPV increases as α decreases.

In the example that tested four predictions of a single hypothesis with the p_FM_ value of 0.005, PPV would be 0.93, i.e. the conclusion should have a 93% chance of being replicated, a marked increase over the 57% chance predicated on a single test α value of 0.05. This agrees with the intuition that conclusions based on multiple lines of evidence should be more secure than those based on one line. This may partially explain why much experimental science appears to be more reliable than the statistical arguments predict.

Investigators could report both the exact probability levels of aggregate value of p_FM_ from Fisher’s Method (or the analogous parameter from a similar test) and the p-value for each experimental result that goes into the aggregate. Reporting both individual and aggregate probabilities would reveal how uniformly the p_FM_ depends on all of the constituent results and help prevent a low p_FM_ that is skewed by one extraordinarily small p-value, from being over-interpreted.

## Discussion

It has been proposed that a significant contributing factor in the reproducibility cricis of science is a widespread tendency for scientists to base major conclusions on the outcomes of single experimental tests. This practice, when combined with low statistical power – generally because of small sample size – has been interpreted to mean that the expected reproducibility of certain sciences, such as neuroscience, will necessarily be poor. The present paper examines underling assumptions of these criticisms and makes three claims: 1) the conclusions of experimental studies emerge from the aggregate outcomes of many tests of a single hypothesis. Therefore, 2) the probability of arriving at such conclusions can be evaluated with Fisher’s Method for Combining Results or a similar procedure. Finally, 3) conclusions that rest on several tests of a hypothesis are more likely to be reproducible than those of large, open-ended studies that are based on a single statistical test.

Although it was not surprising to find that many neuroscience publications reported hypothesis-testing work, the prevalence of this mode (81% of the work) was unexpected. Most papers quickly homed in on one particular idea and tested its implications; moreover, a significant fraction (11/42) of the them referred directly to alternative hypotheses and/or falsifying hypotheses, which are also hallmarks of the hypothesis-testing approach. (Note also that, despite frequent concerns that journals will “not publish negative data,” this evidence shows that they will, at least when it appears in a rigorous, hypothesis-testing context.)

A caveat is that, although the papers came from across the spectrum of neuroscience subfields, they were all published in the *The Journal of Neuroscience* and its Instructions to Authors briefly mentions “hypotheses.” Non-hypothesis-testing papers are plainly not forbidden and there is no discussion of the importance of hypothesis-testing, etc., hence it is unclear how strongly the presence or absence of a hypothesis influences editorial policy, nevertheless it is possible that the papers that I reviewed are not representative of all neuroscience.

One motive for undertaking this inquiry was to try to understand how it is that, if most research findings are “false,” biomedical science can make as much progress as it clearly does make. Part of the answer may be that conclusions that follow from testing hypotheses are more reproducible than the statisticians have recognized. Of course, many factors have been implicated in the reproducibility crisis; indeed, “reproducibility” has a variety of divergent definitions (5) and presumed causes (6). Problems with cell-lines, antibodies, scientific misconduct or other bad behavior stemming from scientific mis-incentives will not be alleviated by purely statistical considerations. Still, statistical arguments are extensively cited in the debate, so it is worth examining them critically.

I chose Fisher’s Method for Combining results as it is simple and well-suited to the context of much current biomedical science. However, Fisher’s Method is one member of a class of similar methods (7), which are often used for meta-analyses, and I have not explored alternatives. It is possible that another method would do as well or better than Fisher’s and this should be looked into in the future. The general comments below should apply to these other methods as well, however.

First, the calculation of p_FM_ can gives a more realistic impression of the strength of a collection experimental results than single-test p-values alone. The use of multiple, focused tests on a single idea would help filter out significant but irreproducible results that can otherwise distract research. The “winner’s curse,” for example, happens when an unusual highly significant result leads to a publication that cannot be duplicated by follow-up studies because the result was basically a statistical aberration (2). “Regression to the mean” dominates the later studies.

Sccond, because Fisher’s Method does not assume a particular significance level for individual test results, its use should help diminish the unhealthy over-emphasis on specific p-values, which can lead to warped priorities and inappropriate incentives for scientists.

Third, Fisher’s Method encourages explicit hypothesis-testing methods which, as we’ve seen, offers greater predicted validity than less well-defined modes. Conducting multiple tests of a hypothesis appears to be a reasonably common practice in neuroscience and having a natural standard method for assessing the reliability of such aggregate results would contribute to effective communication about them. Fisher’s Method demands that the experimental tests be strictly independent of each other and be true predictions of a single hypothesis. “True predictions” are logically valid deductions from the hypothesis, and investigators using the Method would have an incentive to make their reasoning overt. As it stands, ferreting out the implicit hypothesis in a research report and trying to understand how its parts fit together can be a needlessly time-consuming chore. Explicitly stating hypotheses, in and of itself, will undoubtedly improve the clarity of scientific publications and thinking.

Should the aggregate test parameter, p_FM_, have a defined significance level and, if so, what should that be? As shown earlier, p_FM_ will usually be much lower than that the significance level for any single test. While ultimately the probability level that a scientific community recognizes as “significant” will be a matter of convention, it might make sense to require a relatively high level, say ≤ 0.005, for p_FM_. This would mean that, as suggested by the examples above, the average significance of the individual tests would tend to be ~0.05 which, as it is the traditional widely accepted, could foster the adoption of p_FM_ as an additional, informative parameter.

Recently, a large group of eminent statisticians (8) has recommended that science “redefine” its “alpha,” (i.e., significance level) to p < 0.005 from p < 0.05. The lower probability would apply to “new discoveries” but, since the purpose of basic science is to make “new discoveries,” it seems that this standard should apply to virtually all basic science. The standard of p < 0.05 would be reserved for merely “suggestive” evidence. These authors argue that a change in alpha would, among other things, reduce the number of “false positives” that can contribute to irreproducible results with a significance level of p ≤ 0.05. However, another equally large and eminent group of statisticians (9) disagrees with this recommendation, enumerating drawbacks to a much tighter significance level, including an increase in the “false negative rate,” i.e. missing out on a genuine discovery. This second group argues that, rather than changing their alpha level, scientists should take pains to “justify” whatever level they report and recommends doing away with the term “statistical significance” altogether.

I suggest that reserving the more stringent significance level for an aggregate probability value, such as p_FM_, would provide many of the advantages of greater stringency, without the disadvantages of an extremely low p-value. Demanding a more severe significance level for conclusions derived from a number of convergent tests would help screen out the “false positives” resulting from single, atypical test results. On the other hand, a marginal or even presently “insignificant” result, would not be discarded if it were an integral component of a focused group of tests of a hypothesis, which would help guard against the problem of “false negatives.”

Reproducibility is obviously a cardinal virtue for science, however, reproducibility is a huge and complex topic (5,6). and strengthening science will require an accurate assessment of its components. This note argues that some of the statistical concerns about rampant irreproducibility in science are overblown, and suggests that a more realistic interpretation of the scientific process will lead to a clearer picture of the reproducibility issues.

---

**Supplemental Figure 1.**
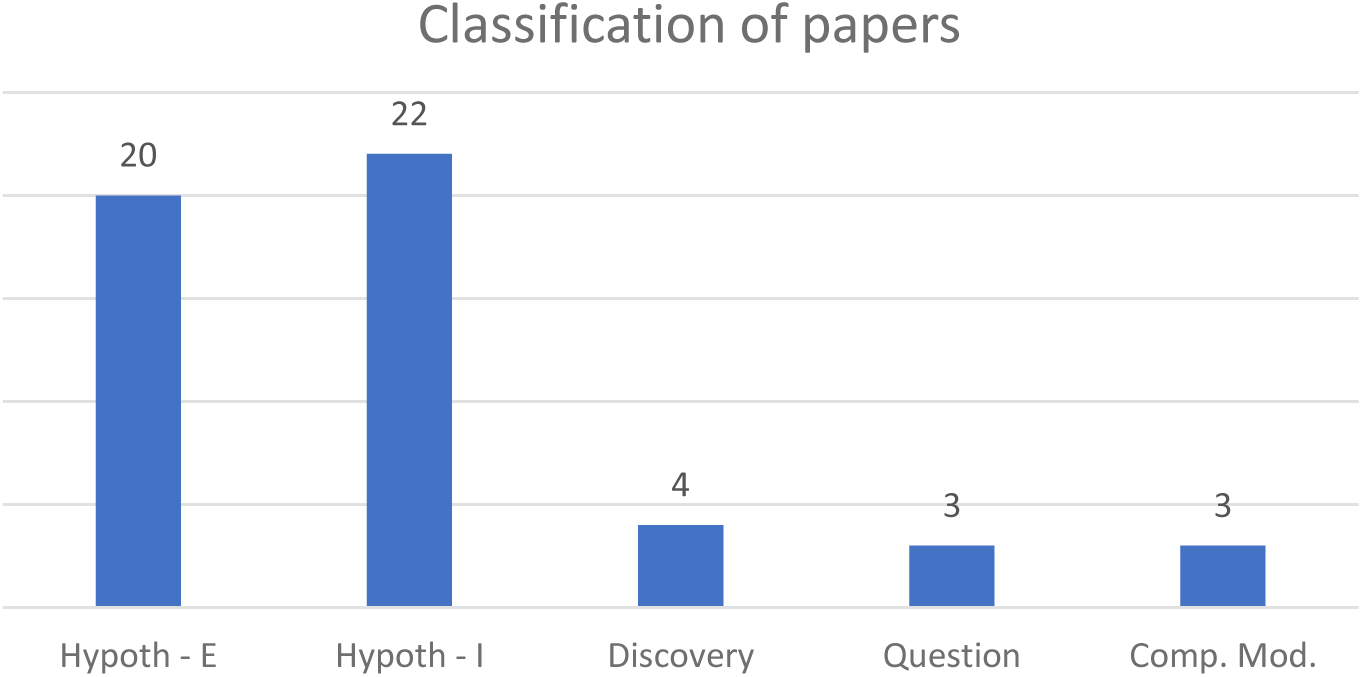
Classification of *Research Articles.* Articles were categorized as based primarily on an explicit hypothesis, implicit hypothesis, “discovery” experiments, open-ended questions, or computer modeling. Numbers in each group above the bars; total n = 52.

**Supplemental Table 1.** Analysis of Journal Articles All *Research Articles* published in the first three issues of *The Journal of Neuroscience* for 2018 (n = 52) identified by Start and End pages. Classifications as follows: Hyp-E – A hypothesis was explicitly identified and a number of its predictions were tested. Hyp-I – A hypothesis was implied but not stated; a number of its predictions were tested. Alt. Hyp. – One or more alternative hypotheses were fairly clearly stated. # Tests – Estimated minimum number of independent tests of the main hypothesis. Support – The experimental tests were all consistent with the main hypothesis. Reject – Some experimental tests explicitly disconfirmed one or more hypotheses. Disc. – Most of the work was purely exploratory. Ques. – Most of the work addressed a series of questions; no evident hypothesis. Comp. – Few experiments, often a great deal of data; mainly computational modeling.

| Start | End | Hyp-E | Hyp-I | Alt. Hyp. | #Tests | Supprt | Reject | Disc. | Ques. | Comp. |
| --- | --- | --- | --- | --- | --- | --- | --- | --- | --- | --- |
| 32 | 50 | X |  |  | 10 | X |  |  |  |  |
| 51 | 59 |  | X | 2 | 8 | X | X |  |  |  |
| 60 | 72 |  | X |  | 7 | X |  |  |  |  |
| 74 | 92 |  | X |  | 7 | X |  |  |  |  |
| 93 | 107 |  | X |  | 7 | X |  |  |  |  |
| 108 | 119 |  |  |  |  |  |  |  | X |  |
| 120 | 136 |  | X |  | 7 | X |  |  |  |  |
| 137 | 148 |  | X | 3 | 8 | X | X |  |  |  |
| 149 | 157 | X |  | 1 | 5 | X | X |  |  |  |
| 158 | 172 |  | X |  | 8 | X |  |  |  |  |
| 173 | 182 | X |  |  | 5 |  | X |  |  |  |
| 183 | 199 | X |  |  | 9 | X |  |  |  | X |
| 200 | 219 |  |  |  |  |  |  | X |  |  |
| 220 | 231 |  |  |  |  |  |  |  | X |  |
| 232 | 244 | X |  |  | 7 | X |  |  |  |  |
| 245 | 256 | X |  |  | 7 | X |  |  |  |  |
| 263 | 277 | X |  | 1 | 7 | X |  |  |  |  |
| 278 | 290 |  | X |  | 8 | X |  |  |  |  |
| 291 | 307 |  | X |  | 10 | X |  |  |  | X |
| 308 | 322 |  | X | 1 | 7 | X |  |  |  |  |
| 322 | 334 |  | X |  | 7 | X |  |  |  |  |
| 335 | 346 |  | X |  | 8 | X |  |  |  |  |
| 347 | 362 |  |  |  |  |  |  | X |  |  |
| 363 | 378 |  | X |  | 11 | X |  |  |  |  |
| 379 | 397 | X |  | 1 | 8 | X | X |  |  |  |
| 395 | 408 |  |  |  |  |  |  |  |  | X |
| 409 | 422 |  | X | 1 | 4 | X |  |  |  |  |
| 423 | 440 | X |  |  | 12 | X |  |  |  |  |
| 441 | 451 | X |  |  | 9 | X |  |  |  |  |
| 452 | 464 |  | X |  | 8 | X |  |  |  |  |
| 465 | 473 | X |  |  | 8 | X |  |  |  |  |
| 474 | 483 | X |  |  | 9 | X |  |  |  |  |
| 484 | 497 | X |  |  | 9 | X |  |  |  |  |
| 498 | 502 |  |  |  |  |  |  |  | X |  |
| 518 | 529 |  | X |  | 8 | X |  |  |  | X |
| 530 | 543 |  |  |  |  |  |  |  |  | X |
| 548 | 554 |  | X |  | 12 | X |  |  |  |  |
| 555 | 574 |  | X |  | 9 | X |  |  |  |  |
| 575 | 585 | X |  |  | 8 |  | X |  |  |  |
| 586 | 594 |  |  |  |  |  |  | X |  |  |
| 595 | 612 | X |  | 1 | 3 | X | X |  |  |  |
| 613 | 630 |  | X |  | 7 | X |  |  |  |  |
| 631 | 647 |  | X |  | 8 | X |  |  |  |  |
| 648 | 658 | X |  |  | 6 | X |  |  |  |  |
| 659 | 678 | X |  | 1 | 5 | X |  |  |  |  |
| 679 | 690 | X |  |  | 12 |  | X |  |  |  |
| 691 | 709 | X |  |  | 2 | X |  |  |  |  |
| 710 | 722 |  |  |  |  |  |  | X |  |  |
| 723 | 732 |  | X | 1 | 5 | X | X |  |  |  |
| 733 | 744 |  |  |  |  |  |  |  |  | X |
| 745 | 754 | X |  | 1 | 4 |  |  |  |  |  |
| 755 | 768 |  | X |  | 5 | X |  |  |  |  |

## References

1. Ioannidis JPA (2005) Why most published research findings are false. PLoS Medicine 2: e124.

2. Button KS, Ioannidis JP, Mokrysz C, Nosek BA, Flint J, Robinson ES, Munafò MR (2013) Power failure: why small sample size undermines the reliability of neuroscience. Nature Reviews Neuroscience 14:365–76.

3. Fisher RA, Statistical Methods for Research Workers; Biological Monographs and Manuals Series, Oliver and Boyd, Edinburgh, 1925); Fisher’s Method (Fisher’s Combined Probability Test) in https://en.wikipedia.org/wiki/Fisher%27s_method. See also, (https://brainder.org/) The logic of the Fisher method to combine P values. Posted on 11.May.2012.

4. https://en.wikipedia.org/wiki/Extensions_of_Fisher%27s_method See also, Brainder (https://brainder.org/) Non-Parametric Combination (NPC) for brain imaging Posted on 08.February.2016.

5. Task Force on Reproducibility, Ameorgrican Society for Cell Biology (2014). How can scientists enhance rior in conducting basic research and reporting research results. White Paper, http://www.acsb.org/reproducibility.

6. Landis SC, Amara SG, Asadullah K, Austin CP, Blumenstein R, Bradley FW, et al. (2012). A call for transparent reporting to optimize the predictive value of preclinical research. Nature. 490:187–191.

7. Winkler AM, Webster MA, Brooks JC, Tracey I, Smith SM, Nichols TE (2016). Non-parametric combination and related permutation tests for neuroimaging. Human Brain Mapping. 37:1486–511.

8. Benjamin DJ, Berger JO, Johannesson M, Nosek BA, Wagenmakers E-J, Berk R et al. (2018). Redefine statistical significance. Nature Human Behaviour 2:6–10.

9. Lakens D, Adolfi FG, Albers CJ, Anvari, Apps MAJ, Argamon SE, et al. (2018) Justify your alpha. Nature Human Behaviour 2:168–171.

